# The Negative Effects of Smoking Cigarettes on Morphological and Histological Damages to 18 and 21 Days Mouse Embryo Liver

**DOI:** 10.1101/2024.06.22.600226

**Authors:** Zahra Peyrovi, Soheil Azizi, Seyed Mohammad Hosseini

## Abstract

Smoking cigarettes can lead to morphological and histological changes in several human and animal tissues. This paper describes the effects of CS exposure on the liver, weight, and length of mouse embryos. 42 NMRI adult mice were used for mating. Pregnant female mice were categorized into 3 groups: F (filtered), NF (non-filtered), and C (controlled). Each group included 14 female mice exposed to 12 cigarettes daily; 7 mice for 18 days and 7 mice for 21 days. In addition, only the group C were exposed to ambient air. Seven mouse embryos from each group were euthanized. Their livers were fixed for histological processing. The Livers of groups F & NF as compared with group C revealed changes in cellular architecture, centrilobular veins, inflammatory cells, kupffer cells, cytoplasmic vacuolation, hepatocytes necrosis, and decreased parenchymal cellularity. The average weight of embryo, liver, and CR in F & NF significantly reduced (p<0.05 and p<0.01).

## Introduction

Tobacco use by pregnant women leads to changes in fetal organ development, which often leads to negative consequences in the offspring ‘s life, such as depression and formation of depression associated immune diseases (1-2-3). Approximately 20%–50% of women report smoking during early pregnancy (4), and 50% of pregnant nonsmokers are exposed to secondhand smoke during pregnancy (5-6). Nicotine penetrates through the placenta into the bloodstream of the fetus. It is also found in the amniotic fluid, from which it penetrates through the skin into the fetus (7-8). The clearness of nicotine and cotinine (the main component of nicotine metabolism) increases in pregnant women (9).

This results from an increased blood flow through the liver, and an increase in the enzymatic decomposition of both nicotine and cotinine in the mother (10-11-12) However, the metabolism in the liver of the fetus is slower, which leads to a longer period of nicotine half-life in the organism of the fetus. This is confirmed by higher concentrations of nicotine observed in the tissues of the fetus, compared to the mother (13-14). The effect of nicotine on the fetus leads to the restriction of intrauterine growth (fetal intrauterine dystrophy), irrespective of the term of delivery (15). Low birth weight of the neonate is the result of the effect of nicotine on the structure and function of the placenta, and disturbances in the supply of oxygen and nutrients to the fetus via the placental barrier. Nicotine also activates nicotinic acetylcholine receptors, resulting in the constriction of blood vessels, and consequently, in the reduction of oxygen supply to the organism of the fetus, leading to the impairment of its development (13-16-17). Tobacco smoking by a pregnant woman results in an increase in the amount of carbon dioxide in her body, which also reduces the oxygen supply to the fetus and inhibits its growth (16). In addition, smoking affects the development of the trophoblastic and results in reduced blood diffusion between the mother and fetus (18-19). Disorders in trophoblastic differentiation under the effect of tobacco smoke occur as early as the beginning of the development of the placenta (20-21). The toxicity of nicotine also consists in inducing the release of oxidants (14-22-23). Insulin sensitivity changes in cells and changes in lipid metabolism profile (24-25-26). The pathophysiology of tissue damage associated with cigarette smoke and the oxidant/antioxidant imbalance caused by the release of reactive oxygen species have been widely studied. Tobacco use causes the release of many substances in the body that have the direct potential of forming free radicals and activating inflammatory cells, such as macrophages and neutrophils, which also produce reactive oxygen species ultimately increasing the tissue concentration of harmful substances (6-27). This study aimed at qualitatively evaluating the inflammatory changes that occur in the liver of neonatal animals that were exposed to cigarette smoke during pregnancy.

## Materials and Methods

During all experiments, animals were kept at ambient temperature (21°C ± 2°C) under a 12-h light/dark cycle (starting at 6:00 pm daily), with food and drinking water ad libitum. For breeding, 42 female and 21 male mice, (approximately 25g and 31g each, respectively) all of the NMRI, were used. Mating occurred through placing 2 female mice and 1 male mouse in a breeding box for a 5-day period. Pregnancy in the female mice was confirmed through observation of the vaginal plug and the vaginal smear (6-41). The day of the vaginal plug was considered as day zero of pregnancy. Pregnant female mice were randomly divided into different groups: Group F (filtered group inhaling cigarette smoke with filter), consisting of 14 pregnant female mice that were subjected to inhalation of cigarette smoke 7 mice for 18 days, and the other 7 for 21 days during pregnancy. Group NF (non-filtered group inhaling cigarette smoke without filter), consisted of 14 pregnant female mice that were subjected to inhalation of cigarette smoke without filter, 7 mice for 18 days and the other 7 for 21 days during pregnancy. Group C (control group), consisting of 14 pregnant female mice that were exposed to ambient air, had 7 mice for 18 days and the other 7 for 21 days during pregnancy. (6-28-29-30-31).

### Exposure to cigarette smoke

Pregnant animals ‘ exposure to cigarette smoke occurred during the 18 & 21 days of gestation and was performed according to procedures described by Ahmadnia et al, and Valencia et al (30-31). In an inhalation chamber, the group F mice were exposed to smoke from 12 commercial filtered cigarettes per day, divided into 3 daily exposures. In an inhalation chamber, the group NF mice were exposed to smoke from 12 commercial non-filtered cigarettes per day, divided into 3 daily exposures. At the time of each exposure, a single cigarette (10 mg of tar, 0.8 mg nicotine, and 10 mg of carbon monoxide) was attached to insufflations. Smoke immediately was introduced into the inhalation chamber where it was collected and it remained for 6 minutes. Following that the chamber was exhausted of cigarette smoke with the fan for 1 minute, and this procedure was repeated for each cigarette (6-30-31).

### Euthanasia and tissue collection

Euthanasia of mouse embryo was performed in 18 & 21 days of pregnancy. In total, 42 from groups F, NF, and C were euthanized. The mice were killed by cervical dislocation and the fetuses were removed by cesarean section. After measuring the fetuses ‘ weight, and length of head to toe (CR), animals were perfused with 10% formalin saline solution. The livers were collected and weighed. The first sample was fixed in buffered formalin solution for at least 48 hours to be used in the histological analysis.

### Histological staining and morphometric analysis

After fixation, the liver tissue samples were processed according to routine histological techniques and embedded in paraffin blocks. Paraffin blocks were sectioned using a microtome into 4 μm thick sections, which were placed onto glass slides, and fixed for histological staining. Staining was performed using hematoxylin-eosin. Liver tissue damage was evaluated on parenchyma showing morphological changes. The tissue section was assessed by two pathologists. The average of ten microscopic fields were recorded by the pathologists for each specimen. Histopathological criteria were defined according to a modified scoring system taken from previous studies(28-32-33-34). The morphological parameters scoring were performed as follows (-) and (+) and (++) and (+++) indicating NO or Few <10%, mild between 10-30%, moderate between 31-50%, and marked changes >50% respectively (33-34-35).

## Statistical analysis

All data was presented as mean ± standard error. Statistical analysis was performed using the ANOVA test, mean comparison was performed by LSD among the 3 groups. The software used was SAS 9.1, differences were considered significant at p<0.05 and p<0.01.

## Results

The average weight of 18 & 21 days of embryo, experimental groups F (filtered) and NF (non-filtered) as compared with the group C (controlled) decreased significantly(respectively p<0.05 and p<0.01). The CR 18 & 21 days in F and NF as compared with the C were significantly decreased (respectively p<0.05 and p<0.01). The average weight of the liver 18 & 21 days in F and NF as compared to the C were significantly decreased (respectively p<0.05 and p<0.01).(Table1) (Fig1).

**Table 1:** Biometric analysis of C, F, and NF. Results are presented as mean± SEM. C: Control group (18 & 21 days); F: Filtered Group Cigarette Smoke during Pregnancy (18 & 21 days); NF: Non-filtered Group Cigarette Smoke during Pregnancy (18 & 21 days); N=7 per, Significantly different between group by ANOVA and LSD test, unpaired for comparison between the 3 groups. Differences were considered significant at (p<0.05) and (p<0.01).

| Experimental Group | C | F | NF | P |
| --- | --- | --- | --- | --- |
| Weight of 18 days (g) | 1.45±0.1 | 1.31±0.32 | 1.11±0.25 | 0.0001<sup>**</sup> |
| CR (cm) of 18 days | 5.33±0.18 | 4.43±0.72 | 4.45±0.37 | 0.0001<sup>**</sup> |
| Weight of 21 days (g) | 1.79±0.15 | 1.65±0.14 | 1.36±0.11 | 0.0001<sup>**</sup> |
| CR (cm) of 21 days | 5.31±0.33 | 4.98±0.31 | 4.54±0.44 | 0.0001<sup>**</sup> |
| Liver 's Weight of 18 days (g) | 0.1±0.01 | 0.09± 0.01 | 0.07± 0.01 | 0.0002<sup>**</sup> |
| Liver 's Weight of 21 days (g) | 0.15±0.00 | 0.13±0.00 | 0.11±0.00 | 0.0001<sup>**</sup> |

**Figure 1:**
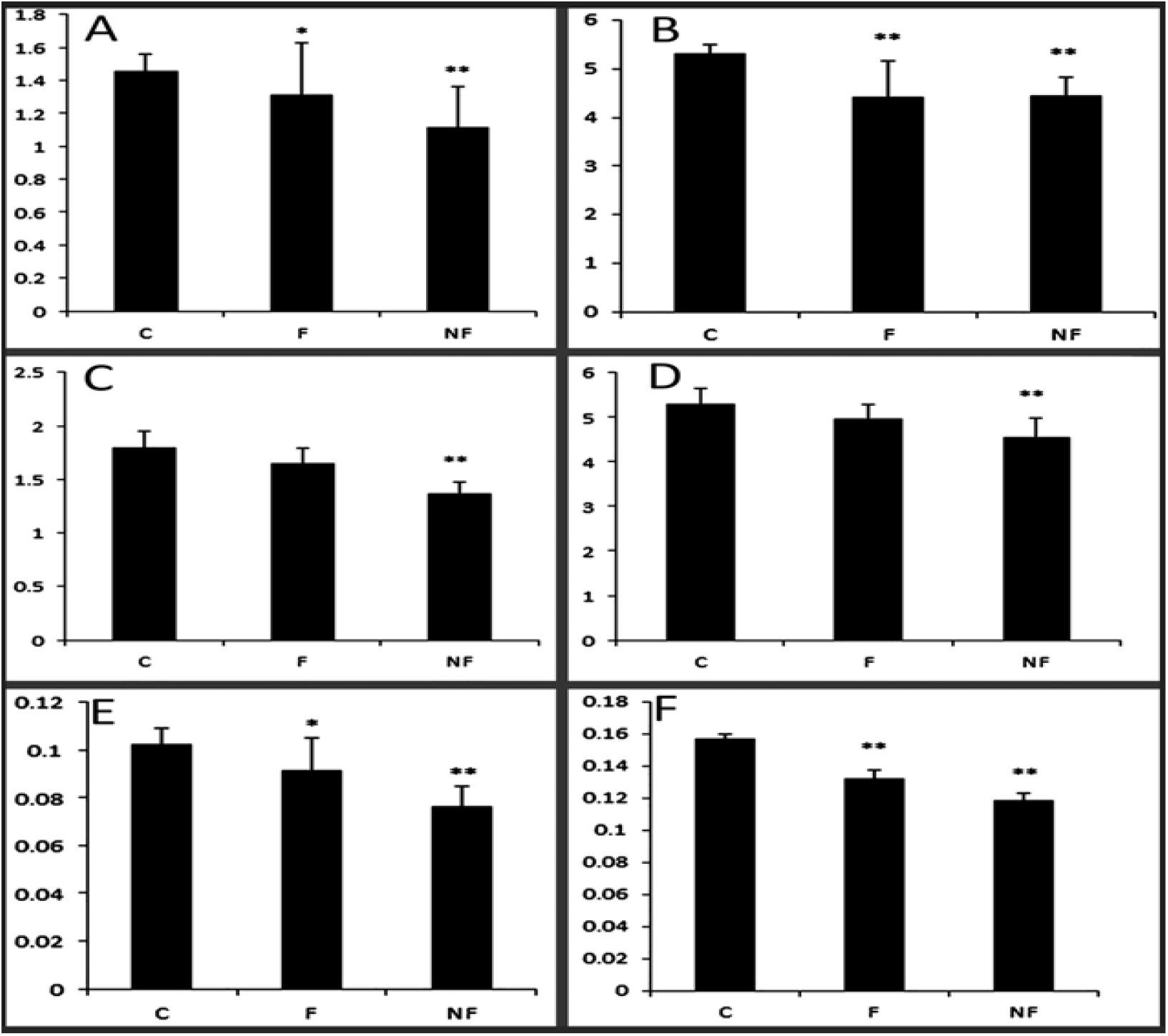
Morphometric weight, CR, and liver of 18 & 21 days mouse embryo analysis. A) The Weight of an 18-day mouse embryo. B) The CR of the 18-day mouse embryo. C) The Weight of the 21-day mouse embryo. D) The CR of the 21-day mouse embryo. E) The liver’s weight of 18 days mouse embryo. F) The liver’s weight of 21 days mouse embryo. Experimental Groups; C: Control Group during Pregnancy; F: Filtered Group Cigarette Smoke During Pregnancy; NF: Non-filtered Group Cigarette Smoke during Pregnancy; 18 & 21 days of pregnancy; N=7 per group.

## Histopathological finding of liver tissues

The results of histopathology changes in the liver tissues are summarized in table (2,3). Histological findings of the control groups 18 & 21 days of embryo revealed normal trabecular no variation in hepatocyte size and with certain cellular limits, normal capillaries form architecture and absent or with few hydropic degenerations, cytoplasmic vacuolation, hepatocytes necrosis, hepatocytes poly nuclear, infiltration of inflammatory cells, activated kupffer cells (Fig 2 (arrow 2,5)). Section of the liver of group F 18 & 21 days and group NF 18 & 21 days show mild and moderate changes in cellular architecture, congestion and dilation of centrilobular veins and sinusoids, inflammatory cells infiltration, activated kupffer cells, cytoplasmic vacuolation, hepatocytes necrosis, hepatocytes poly nuclear and parenchymal decreased cellularity, respectively (Fig 2 (arrow 3,4,6,7)). Moderate and marked hydropic degeneration in F and NF groups are also seen, respectively.

**Table 2:**
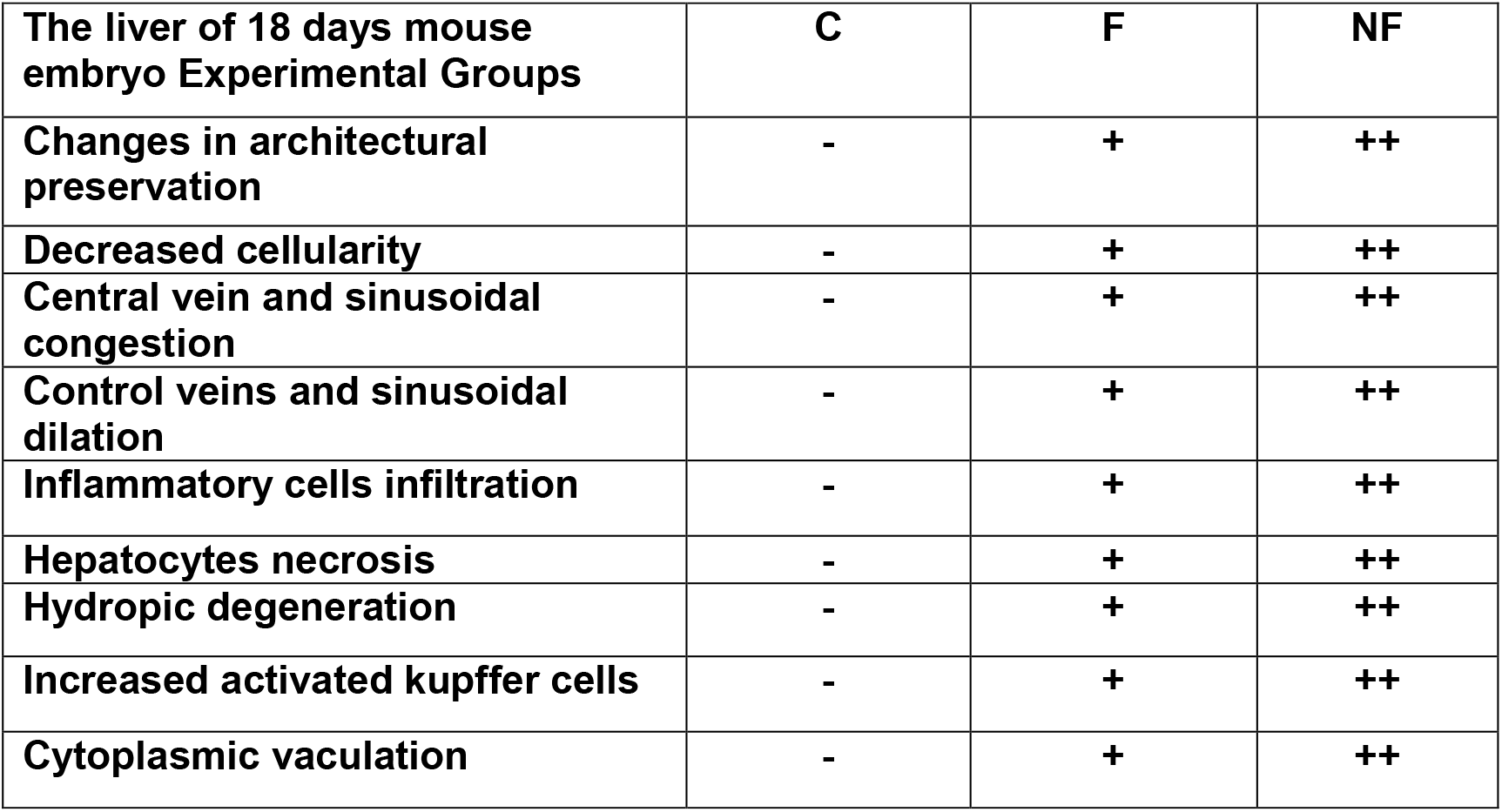
Histopathological changes in 3 groups, C: Control Group during Pregnancy; F: Filtered Group Cigarette Smoke during Pregnancy; NF: Non-filtered Group Cigarette Smoke during Pregnancy; The liver of 18 days mouse embryo.

| **The liver of 18 days mouse embryo Experimental Groups** | **C** | **F** | **NF** |
| --- | --- | --- | --- |
| **Changes in architectural preservation** | - | + | ++ |
| **Decreased cellularity** | - | + | ++ |
| **Central vein and sinusoidal congestion** | - | + | ++ |
| **Control veins and sinusoidal dilation** | - | + | ++ |
| **Inflammatory cells infiltration** | - | + | ++ |
| **Hepatocytes necrosis** | - | + | ++ |
| **Hydropic degeneration** | - | + | ++ |
| **Increased activated kupffer cells** | - | + | ++ |
| **Cytoplasmic vacuolation** | - | + | ++ |

**Table 3:** Histopathological changes in 3 groups, C: Control Group during Pregnancy; F: Filtered Group Cigarette Smoke during Pregnancy; NF: Non-filtered Group Cigarette Smoke during Pregnancy; The liver of 21 days mouse embryo.

| **The liver of 21 days mouse embryo Experimental Groups** | **C** | **F** | **NF** |
| --- | --- | --- | --- |
| **Changes in architectural preservation** | - | + | ++ |
| **Decreased cellularity** | - | + | ++ |
| **Central vein and sinusoidal congestion** | - | + | ++ |
| **Control veins and sinusoidal dilation** | - | + | ++ |
| **Inflammatory cells infiltration** | - | + | ++ |
| **Hepatocytes necrosis** | - | + | ++ |
| **Hydropic degeneration** | - | + | ++ |
| **Increased activated kupffer cells** | - | + | ++ |
| **Cytoplasmic vaculation** | - | + | ++ |

**Figure 2:**
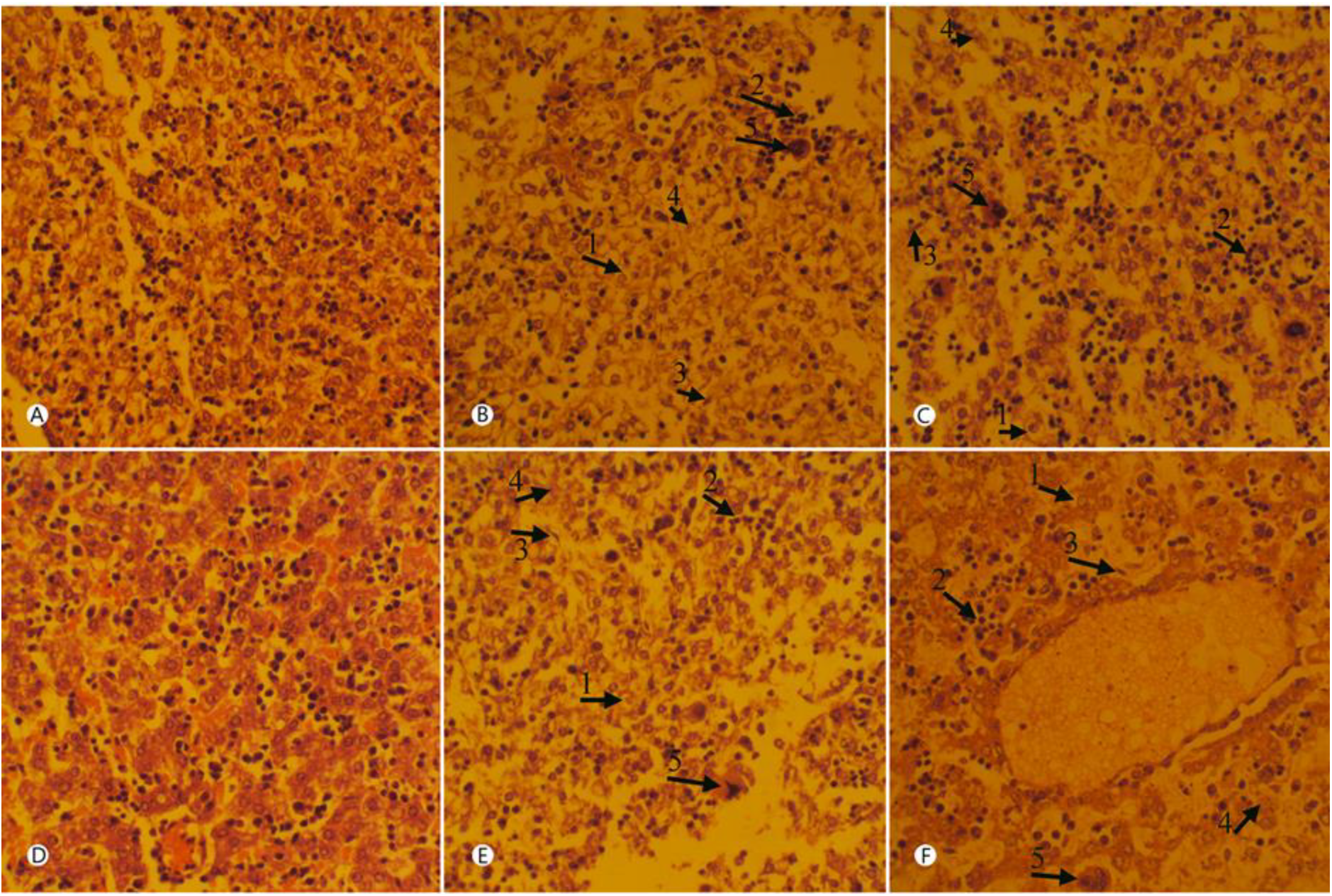
Histological section of liver of 18 & 21 days mouse embryo; A) Group C of 18 days; Normal tissue, H&E, X40. B) Group F of 18 days; C) Group NF of 18 days; In Group F & NF of 18 days: (arrow 1) indicates Hydropic degeneration, (arrow 2) indicates Inflammatory cells infiltration, (arrow 3) indicates Increased kupffer cells, (arrow 4) indicate Hepatocytes necrosis, (arrow 5) indicate hepatocyte poly nuclear, H&E, X40. D) Group C of 21 days; Normal tissue, H&E, X40. E) Group F of 21 days; F) Group NF of 21 days; In Group F & NF of 21 days: (arrow 1) indicates Hydropic degeneration, (arrow 2) indicates Inflammatory cells infiltration, (arrow 3) indicates Increased kupffer cells, (arrow 4) indicate Hepatocytes necrosis, (arrow 5) indicate hepatocyte poly nuclear, H&E, X40.

## Discussion

Results indicated that a decrease in the average weight of embryo and CR 18 & 21 days in the F & NF, filtered and non-filtered groups were more than in the C, controlled group, the difference was significant (6-36-37-38-39). The average weight of the liver at 18 & 21 days in the F & NF, filtered, and non-filtered groups as compared with the C control group were significantly decreased (6-39). Histological findings in our study like Diniz Mf et al(6), which is a similar model exposure to cigarette smoke during pregnancy cause pathological changes that indicate infiltration of inflammatory cells and microvascular steatosis that were not seen in the controlled group. According to Hanene et al (39), and Samuel Santos Valença et al (40), relatively like models effect of the smoke of tobacco nicotine, oral nicotine, and subcutaneous injection of nicotine on histological changes in the rat ‘s liver that caused loss of trabecular arrangement, decreased cellularity, congestion, degeneration and necrosis in our study are also seen.

## Conclusion

Our studies examined changes in arrangement (6-29-39) and histopathological parameters. In future studies consisting of biochemical, immunological, and molecular pathology, further research will be necessary to determine the mechanisms responsible for these changes in smoking cigarettes.

## Notes

### Competing Interest Statement

The authors have declared no competing interest.

